## Supplementary Note for "Integrated multiplexed assays of variant effect reveal *cis*-regulatory determinants of catechol-*O*-methyltransferase gene expression"

Untranslated regions influence variant effects on MB-COMT transgene expression

Our analysis of ribosome profiling data for our MB-COMT transgene and endogenous TIS sites indicated uORFs in the 5′ UTR of *COMT* (**Figure 1C, Supplementary Figure 1B**). These uORFs are 74 and 28 nt upstream of the start codon for MB-COMT and are predicted to generate short peptides of 23 and 69 amino acids, respectively. However, because of our inclusion of a N-terminal Flag tag, uORF B no longer significantly overlaps the canonical MB-COMT ORF due to translation termination within the tag. To test whether these uORFs alter expression of COMT, we generated variants at the uORF A and B start codons (CUG>UUG); generated a silent variant in Flag that restores the endogenous uORF B frame; and deleted the Flag tag. Although we found no impact on protein abundance with the uORF A start codon variant, we found the uORF B start codon variant and Flag tag variants increased COMT protein abundance two to three-fold (two-sided Wilcoxon rank sum test p-value $< 2.2 \times 10^{-16}$, **Supplementary Figure 1C-E)**, suggesting uORF B represses translation of canonical MB-COMT. Additionally, deletion of poly-rC-binding protein (PCBP) motifs in the 5′ UTR led to a slight but significant increase in COMT protein abundance (~36% increase, Wilcoxon p-value $< 2.2 \times 10^{-16}$).

We serendipitously isolated a 53 nt deletion in the 5′ UTR of *COMT* during cloning that encompasses the putative uORFs (**Supplementary Figure 2A**). We assayed *COMT* mRNA abundance for the 5′ UTR deletion and 5′ UTR-containing control, and found the 5′ UTR deletion exhibited two-fold lower *COMT* mRNA abundance (**Supplementary Figure 2B**). We also ran polysome profiling experiments and determined that both the 5′ UTR-containing and 5′ UTR deletion mRNAs were highly loaded onto polysomes, in agreement with previous polysome profiling data generated for the endogenous *COMT* gene (Floor and Doudna, 2016). Consistent with RT-qPCR readout, the 5′ UTR deletion showed reduced mRNA abundance in ribosomal subunits and the monosome to polysome fractions. Yet, we found no significant global shift in ribosome loading across polysomal fractions (**Supplementary Figure 2C**).

In contrast, the endogenous 5′ UTR had an inhibitory effect on protein abundance. By flow cytometry, the 5′ UTR deletion exhibited increased COMT protein abundance (**Supplementary Figure 2D**, two-fold increase in median fluorescence). Yet, mCherry fluorescence readout was similar between the 5′ UTR deletion and control, despite a modest difference reported by RT-qPCR. We suggest the discrepancy between mRNA abundance measurements by flow cytometry and RT-qPCR may be due to the long half-life of the mCherry protein (Matreyek et al., 2020). Yet, in agreement with flow cytometry results, Flag-immunoblotting reported 78% higher abundance for the 5′ UTR deletion (**Supplementary Figure 3C**).

We conclude that an element, or multiple elements, in the *COMT* 5′ UTR represses MB-COMT translation and reduces protein abundance. Our results are consistent with a mechanism in which uORF translation termination near the canonical start codon represses translation of MB-COMT within our transgene (**Supplementary Figure 2E, Limitations**). An alternative interpretation is that disruption of RNA secondary structure near the canonical TIS increases the rate of initiation (Tsao et al., 2011).

<u>Common population variants have no effect on MB-COMT transgene expression</u>
Prior characterization of haplotypes comprised of the coding SNPs rs4680, rs4818, and rs4633, which form low-, average-, and high-pain sensitivity phenotypes (LPS, APS, HPS), indicated APS and HPS haplotypes decrease enzyme activity compared to the LPS haplotype (Nackley et al., 2006). However, these effects were only observed when the haplotypes were expressed from a full-length *COMT* transcript with 5′ and 3′ UTRs (Nackley et al., 2006) (Supplementary Figure 6). In addition, the APS haplotype exhibited *increased* protein abundance in an *in vitro* context and in HEK293 and MCF-7 cells, but not COS-1 or HepG2 cells. Increased protein abundance for the APS haplotype was hypothesized to result from a less stable secondary structure at the TIS of S-COMT owing to the rs4633 C>T variant. These results suggested that expression phenotypes of the common haplotypes in *COMT* are cell-type specific and depend on either the 5′ UTR, 3′ UTR, or both UTRs.

We individually assayed five common population variants for effects on *COMT* RNA and protein abundance (LPS haplotype without rs6269): rs4680 (G>A), rs4818 (G>C), rs4633 (C>T); as well as rs6267 (G>T) and rs74745580 (C>T), which were previously associated with disease phenotypes or reported to alter *COMT* expression (Gothelf et al., 2014; Lin et al., 2017; Li et al., 2014) (**Supplementary Figure 3A**). These single variants enabled us to test effects for the partial HPS (rs4818) and LPS-T166 (rs4633) haplotypes (lacking rs6269), which led to low and high protein abundance, respectively (Tsao et al., 2011). In our transgene, we found no difference in mRNA or protein abundance for any of the variants by flow cytometry or immunoblotting (**Supplementary Figure 3B-C**). Our results suggest that the prior observed effects of the common population variants may require expression from a native context (including endogenous UTRs).

Importantly, our study does not rule out a model where rs4633 facilitates translation initiation. Nevertheless, our data suggest a potential concurrent mechanism where rs4633 leads to higher protein abundance in human cell lines and in an *in vitro* translation assay (Tsao et al. 2011) by increasing RNA abundance. We note that Tsao et al did not directly measure RNA abundance in their study. In Supplementary Figure 3A of Nackley et al 2006, the APS haplotype containing rs4633 C>T showed slightly higher total RNA abundance compared to the LPS haplotype (in our study, the wild-type template). However, this was not statistically significant and was only observed for the S-COMT isoform. It is possible that our observations are compatible with the conclusions in Tsao et al. 2011. For example, increased translation of rs4633 C>T may lead to stabilization of the RNA.
